# Deletion of mouse *Setd4* promotes the recovery of hematopoietic failure

**DOI:** 10.1101/860239

**Authors:** Xing Feng, Huimei Lu, Jingyin Yue, Megha Shettigar, Jingmei Liu, Lisa K Denzin, Zhiyuan Shen

**Affiliations:** Rutgers Cancer Institute of New Jersey, Department of Radiation Oncology, Rutgers Robert Wood Johnson Medical School, 195 Little Albany Street, New Brunswick, New Jersey 08901, USA; Child Health Institute of New Jersey, Rutgers Robert Wood Johnson Medical School, Rutgers University, New Brunswick, New Jersey, USA

**Keywords:** SETD4, radiation-induced bone marrow failure, hematopoiesis, hematopoietic stem cell (HSC), bone marrow niche

## Abstract

Acquired hematopoietic failure is commonly caused by therapeutic and accidental exposure to toxic agents to the bone marrow (BM). Efficient recovery from the BM failure is not only dictated by the intrinsic sensitivity and proliferation capacity of the hematopoietic stem and progenitor cells, but also nourished by the BM environment niche. Identification of genetic factors that improve the recovery from hematopoietic failure is essential. Vertebrate SETD4 is a poorly characterized, putative non-histone methyl-transferase whose physiological substrates have not yet been fully identified. By inducing *Setd4* deletion in adult mice, we found that loss of *Setd4* improved the survival of whole body irradiation induced BM failure. This was associated with improved recoveries of long-term and short-term hematopoietic stem cells (HSC), and early progenitor cells. BM transplantation analyses surprisingly showed that the improved recovery was not due to a radiation resistance of the *Setd4* deficient HSC, but that *Setd4* deficient HSC were actually more sensitive to radiation. However, the *Setd4* deficient mice were better recipients for allogeneic HSC transplantation. Furthermore, there was an enhanced splenic erythropoiesis in *Setd4* deficient mice. These findings not only revealed a previously unrecognized role of the *Setd4* as a unique modulator of hematopoiesis, but also underscored the critical role of the BM niche in the recovery of hematopoietic failure. These studies also implicated *Setd4* as a potential target for therapeutic inhibition to improve the conditioning of the BM niche prior to allogeneic transplantation.

**Key points:**

- Deletion of *Setd4* in adult mice improved the survival from hematopoietic failure.
- *Setd4* deficiency sensitized HSCs to radiation, but improved bone marrow environment niche.
- The study suggests that SETD4 as a potential inhibitory target to improve bone marrow niche function for recovery of bone marrow failure.

## Introduction

The mammalian hematopoietic system is subject to a consistent renewal during the life span of an individual. Inherited and acquired hematopoietic failure can be caused by diseases such as myelodysplastic syndrome, acute myeloid leukemia, and Fanconi anemia(1-3), as well as a consequence of chemotherapy treatment or accidental over-dose exposure to radiation and other toxic agents(4-7). Depleting the host bone marrow (BM) hematopoietic cells before allogeneic hematopoietic stem cell (HSC) transplantation is commonly used to treat a wide array of malignancies including multiple myeloma, leukemias, lymphomas, and some solid tumors(8-10). Understanding the molecular factors that modulate hematopoietic recovery is essential to improve the outcomes of these diseases and treatments.

Lysine methylation plays an important role in diverse cellular processes to regulate chromatin structures, genome stability, and fate determination of stem cells, etc (11-19). Many lysine methyl-transferases harbor a Su(var)3-9-Enhancer of zeste–Trithorax (SET) domain, which is responsible for transferring a methyl group from S-adenosylmethionine to lysine residues. The human genome encodes an estimated of 58 SET domain methyltransferases(20). Although most of the SET domain proteins have documented histone modifying activity, the SETD6 group of SET proteins (including SETD6, SETD3 and SETD4) represents a distinct class of non-histone methyltransferases(20), and they share a similar substrate-binding domain resembling that of the plant Rubisco methyltransferase(21-27). SETD6 methylates p65 (RelA)(24,28), and other non-histone proteins(29-31). SETD3 was initially reported to methylate histones(32,33), but recently found to a histidine residue in β-actin(25,26,34). However, little is known about the function of mammalian SETD4.

The brine shrimp *Artemia Parthenogenetica* SETD4 was initially suggested to play a role in methylating histone H4K20 and H3K79(21), and the human SETD4 protein may have the same activity (35). However, another report failed to identify these sites as SETD4 substrates, and instead suggested H3K4 as the substrate of SETD4 in macrophages (36). Emerging evidence suggests that SETD4 may have an oncogenic activity(37-39). According to the TCGA database, oncogenic fusion of SETD4 was found to be with TMPRSS2 in prostate cancer, with FTCD in invasive ductal carcinoma, with KIAA1958 in serous ovarian cancer, and with B4Galt6 in lung squamous cancer. SETD4 was highly expressed in the quiescent breast cancer stem cell sub-population of MCF7 cells(35), and an induced *Setd4* deletion in adult mice delayed the radiation-induced T-lymphoma development in mice(40).

In this study, we found that induced *Setd4* knockout in adult mice improved the hematopoietic recovery after total body irradiation (TBI). Interestingly, the improved recovery was not associated with a resistance of the *Setd4* deficient HSCs to radiation, rather the *Setd4* deficient HSC were actually more sensitive to radiation than wild type HSC *in vivo*. However, *Setd4* deficient mice were more receptive to HSC transplantation from wild type donors, and that *Setd4* deficient HSCs had a repopulating advantage in recipient mice after transplantation. These results underscore the unique mode by which *Setd4* regulates hematopoiesis and implied that Setd4 inhibition may not only improve the competence of donor HSCs but also the ability of the recipient BM to receive the transplantation.

## Materials and Methods

*Detailed materials and methods can be found in the Supplemental Materials and Methods.*

### Mouse lines, tamoxifen (Tam) treatment, and TBI

The sources and properties of three mouse lines (*Setd4*^*flox/flox*^, *Rosa26-CreERT2*^*/+*^, *Tp53*^*flox/flox*^) used in this study are summarized in Table S1. The construction, verification, and genotyping strategies of a mouse line with an allele of floxed exon-6 of Setd6, designated Setd4^flox/flox^, were described in details(40). Crossing of the Setd4^flox/flox^ with *Rosa26-CreERT2* (B6.129, Jackson Laboratory Stock No 008463) produced the Setd4^flox/flox^;Rosa26-CreERT2 mice. Adult Setd4^flox/flox^;Rosa26-CreERT2 mice were i.p. injected with Tam (or corn oil as the control) to induce *Setd4* deletion(40). Mice were temporarily restrained in a disk-shaped container and whole-body irradiated with γ-rays.

### Analysis of BM and peripheral blood cells

BM single-cell suspensions were counted to determine the total number of cells. Then, aliquots were stained for surface markers to determine the relative abundance of each cell types(41). The sources of antibodies are listed in Tables S2 and S3. For BM transplantation, a given number of sorted donor Lin-cells were transplanted into lethally irradiated recipient mice. Blood was collected to monitor the success of the transplantation based on the donor CD45.1 or CD45.2 congenic markers.

### Statistics

Statistical analyses were performed using Prism GraphPad, including Student’s *t*-test and Log-rank (Mantel-Cox) test.

## Results

### Deletion of *Setd4* protects mice from ionizing radiation (IR) induced BM failure

To avoid any potential impact of a constitutive *Setd4* deletion on the embryonic and early postnatal development of mice, the Tam-inducible knockout mouse model (*Setd4* ^*flox/flox*^;*Rosa26-CreERT2*^*+*^) (40) was used in this study. Whenever possible, sex-matched littermates were treated with Tam (*Setd4*^*Δ*^) or Oil (*Setd4*^*flox*^). We challenged these mice with TBI of 8 or 9 Gy that cause approximately 50% or 90% death due to hematopoietic failure for this strain of mice. Surprisingly, we found that *Setd4*^*Δ/Δ*^ mice survived significantly better than the mice treated with Oil when exposed to 8 Gy TBI (Fig. 1A). When the mice were exposed to a higher dose of 9 Gy (Fig. 1B), the *Setd4*^*Δ/Δ*^ mice survived significantly better than the control mice, but the *Setd4*^*Δ/wt*^ mice had similar survival rate as the control mice. Tam-treatment of wild type mice (including *Setd4*^*flox/flox*^;*Rosa26CreERT2*^*-/-*^, *Setd4*^*flox/wt*^;*Rosa26CreERT2*^*-/-*^, and *Setd4*^*wt/wt*^;*Rosa26-CreERT2*^*+*^) did not alter the overall survival of the mice (Fig. 1C). It is interesting that the improved survival of the *Setd4*^*Δ/Δ*^ mice was similar to the level of *p53*^*Δ/Δ*^ mice, but co-deletion of *p53* and *Setd4* had an additive effect and better protected the mice from irradiation (Fig. S1).

**Figure 1.**
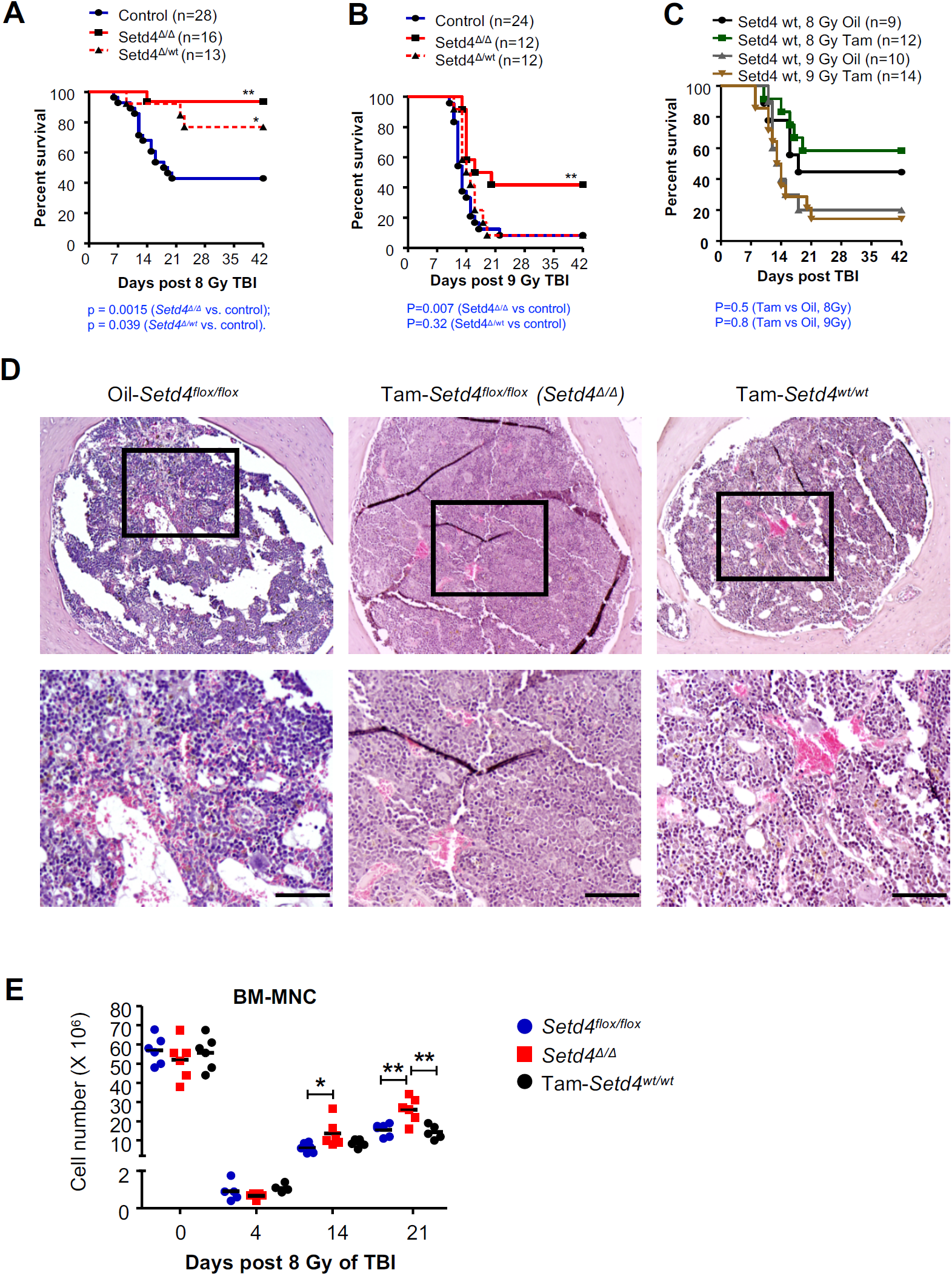
Sensitivity of Setd4 deficient mice to TBI. (A&B) Tam or control treated sex-matched littermates of *Setd4*^*flox/flox*^;*Rosa26-CreERT2*^*+*^ or *Setd4*^*flox/wt*^;*Rosa26-CreERT2*^*+*^ were divided into three experiments groups: 1) Control: Oil-treated *Setd4*^*flox/flox*^;*Rosa26-CreERT2*^*+*^ or *Setd4*^*flox/wt*^;*Rosa26-CreERT2*^*+*^; 2) *Setd4*^*Δ/wt*^: Tam-treated *Setd4*^*flox/wt*^;*Rosa26-CreERT2*^*+*^; and 3) *Setd4*^*Δ/Δ*^: Tam-treated *Setd4*^*flox/flox*^;*Rosa26-CreERT2*^*+*^. Mice were treated with 8 Gy (A) or 9 Gy (B) of TBI, and monitored for survival. The P-values and number of mice in each group are shown. (C) Control mice treated with tamoxifen and Oil and then TBI as indicated. Oil: Oil-treated *Setd4*^*flox/flox*^;*Rosa26CreERT2*^*-/-*^, *Setd4*^*flox/wt*^;*Rosa26CreERT2*^*-/-*^, and *Setd4*^*wt/wt*^;*Rosa26-CreERT2*^*+*^ mice. For 8Gy, 4, 3 and 2 mice of the above genotypes were included. For 9 Gy, 6, 2, and 2 mice were used. Tam: Tam-treated wt mice with the same genotypes above. For 8Gy, 4, 4, and 4 mice with the same three genotypes were used. For 9 Gy, 6, 4, and 4 were included. (D) Repopulation of BM cells after TBI. Hematoxylin-and-eosin (HE) representative images from the staining of cross sections of femur BM cavities from the indicated group of mice 14 days after 8 Gy of TBI. Hematopoietic cells clusters had dramatically repopulated the bone marrows of Tam-treated *Setd4*^*flox/flox*^;*Rosa26CreERT2*^*+*^ (*Setd4*^*Δ/Δ*^) mice, but not in Oil-treated *Setd4*^*flox/flox*^;*Rosa26CreERT2*^*+*^ (*Setd4*^*flox/flox*^) or the Tam-treated *Setd4*^*wt/wt*^;*Rosa26-CreERT2*^*+*^ *(Tam-Setd4*^*wt/wt*^). Upper panel, 40x; lower panel, 100x; scale bar = 100μm. (E) The total number of BM mono-nuclear cell (BM-MNC) collected from a one paired femur and tibia.

It is well established in the radiation biology field that radiation-induced BM failure often results in death within 30 days and mostly between 2-3 weeks. The results in Fig. 1A-B suggested that *Setd4* deficient mice had a decreased sensitivity to radiation induced BM failure. To verify this, we performed Hematoxylin-and-Eosin (HE) staining of the femur BM cavities 14 days after 8 Gy TBI. We designed our approaches with three animal groups: 1) Oil-treated *Setd4*^*flox/flox*^;*Rosa26-CreERT2*^*+*^ (*Setd4*^*flox/flox*^) littermate mice as a control; 2) *Setd4*^*Δ/Δ*^; and 3) Tam-treated *Setd4*^*wt/wt*^;*Rosa26-CreERT2*^*+*^ (Tam-*Setd4*^*wt/wt*^) as another control. We found that hematopoietic cells clusters had dramatically repopulated the BM cavity of *Setd4*^*Δ/Δ*^ mice but not those of *Setd4*^*flox/flox*^ and Tam-*Setd4*^*wt/wt*^ mice (Fig. 1D). Furthermore, we isolated the BM cells and determined the number of the BM mono-nuclear cells (BM-MNC) in the survived mice. Four days after irradiation, there were equal depletion of BM-MNC for knockout and the two control groups (Fig. 1E). However, by day 14 and 21, there was more BM-MNC in the *Setd4*^*Δ/Δ*^ mice than control mice, suggesting an enhanced recovery of BM-MNC when *Setd4* was deleted.

The mice that survived the first 30 days from the hematopoietic syndrome after 8 Gy of TBI were kept for long-term observation. No major differences in term of overall survival (Fig. S2), or in the incidences of tumors were observed. This result suggests that once the mice survived the initial hematopoietic failure, Setd4 loss did not significantly impact the overall survival or tumor development. Furthermore, we irradiated the littermates of *Setd4*^*Δ/Δ*^ and *Setd4*^*flox/flox*^ mice with 13 Gy TBI, and all mice died within 10 days post TBI (Fig. S3), which is typical of radiation induced GI syndrome. The similar sensitivity of these mice in both groups suggested that there was no effect of *Setd4* loss in protecting the mice from radiation induced GI syndrome.

### *Setd4* deletion promotes BM recovery

Since the deletion of *Setd4* protected mice from the fatality of radiation induced hematopoietic syndrome, we hypothesized that *Setd4* may modulate BM recovery from radiation damage. To test this hypothesis, we quantified the number of different cell types in the hierarchy of the BM regeneration with well-established biomarkers and methods(41-44) (detailed in Fig S4 and Supplement Materials and Methods). Representative flow cytometry profiles for a pair of littermates of *Setd4*^*Δ/Δ*^ and *Setd4*^*flox/flox*^ are shown in Fig. S4B.

The lineage relationships of these cell populations are simplified in Fig. 2A. Among the LSK population of cells (Fig. 2B-F), which represent the stem and early progenitor cells, there were no significant difference of total LSK, LT-HSC, ST-HSC, MPP, and LMPP populations between *Setd4* knockout mice and control mice without irradiation. Shortly after irradiation, all cell populations were equally depleted at 4 days. However, by 2 weeks after irradiation, the *Setd4* knockout mice had an enhanced recovery of LT-HSC, ST-HSC, and MPP, but not LMPP populations. Among the LS^−^K and the LS^low^K^low^ populations, which contains the committed progenitor cell populations (Fig. 2G-J), there were enhanced recoveries of the CMP, MEP, and the GMP populations by 2 weeks, but not the CLP population. By 3 weeks post TBI, the difference among these groups were largely diminished, which is expected because only the mice that survived were available for this analysis and these mice had survived past the critical time period of hematopoietic failure and had likely succeeded in replenishing damaged BM at 3 weeks post TBI. Together, these data indicate that *Setd4* inactivation leads to an improved recovery of IR-induced BM injuries by more efficient expansion of early stem and progenitor cells, such as LT-HSC, ST-HSC, and the myeloid progenitors, but not the LMPP and CLP which ultimately repopulate the peripheral lymphocyte pools.

**Figure 2.**
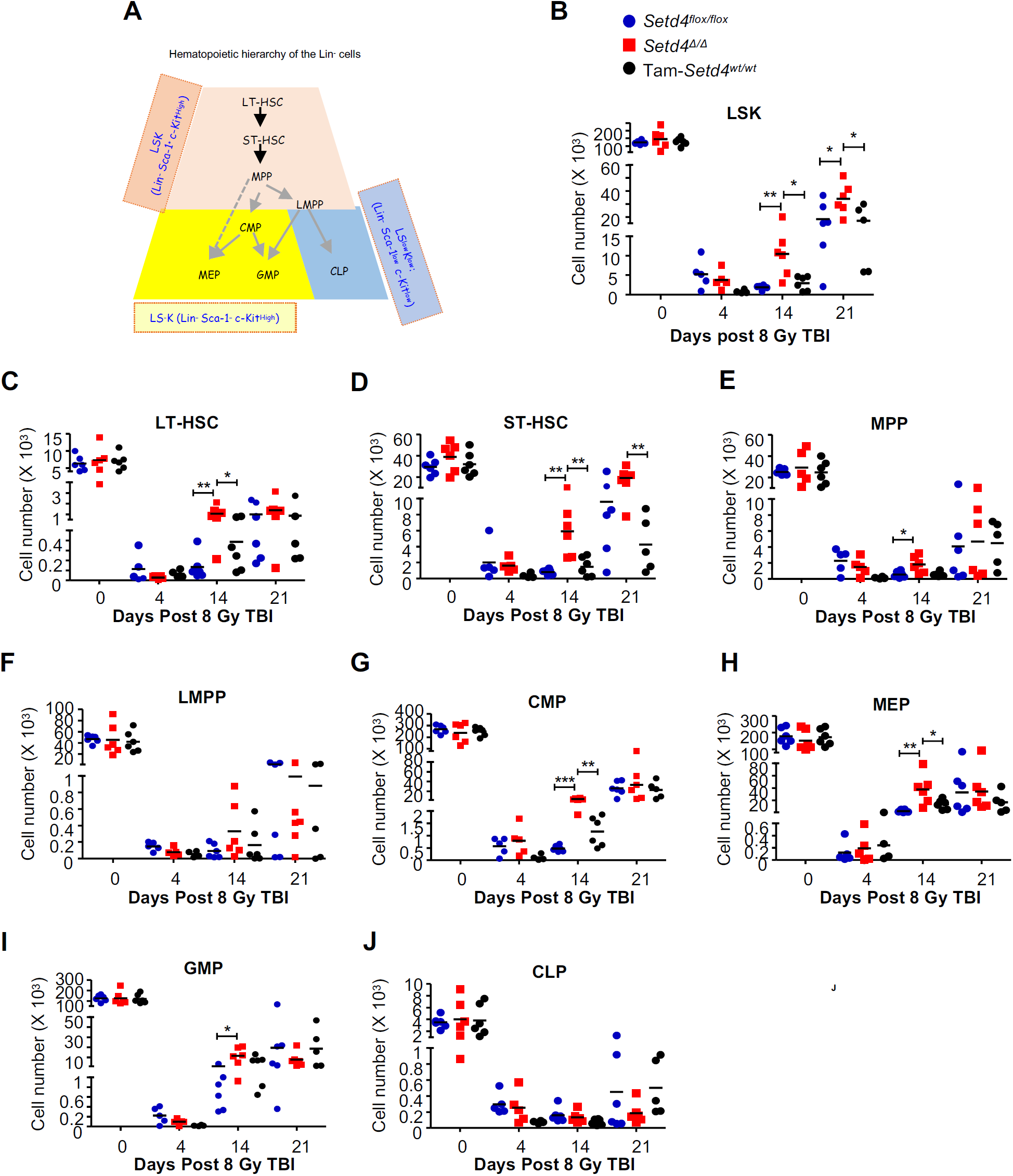
Cell counts of hematopoietic stem cells (HSCs) and progenitor cells in the BM after 8 Gy of irradiation. (A) A simplified illustration of the lineage hierarchy of the Lin^−^ cells. Based on Sca-1 and c-Kit expression, the Lin- cells were divided into three populations shown with different colors. In each population, the specific cell types can be identified by additional cell markers (see Supplemental Materials and Methods, and Fig. S4 for details). LSK: Sca-1^+^ and c-Kit^+^; LS^−^ K: Sca-1^−^ and c-Kit^+^; LS^low^K^low^: Sca-1^low^ c-Kit^low^; LT-HSC: long-term HSC; ST-HSC: short-term HSC; MPP: multipotent progenitor; LMPP: lymphoid-primed MPPs; CMP: common myeloid progenitors; GMP: granulocyte-macrophage progenitors; MEP: megakaryocyte-erythroid progenitors. CLP: common lymphoid progenitors. (B-J) The numbers of total LSK, LT-HSC, ST-HSC, MPP, LMPP, CMP, MEP, GMP, and CLP cells at different the indicated times after 8 Gy of TBI. The BM-MNC (Fig. 1E) were then used to sort out the Lin-cells, and to determine the relative percentage of each of the cell types in the hematopoietic hierarchy as labeled above each panel. The designation of animal genotypes is shown in Panel B. For each tested mouse, a pair of femur and tibia were crushed to collect all BM cells at 4, 14, and 21 days after 8Gy of TBI. The mouse genotypes were: 1) Oil-treated *Setd4*^*flox/flox*^;*Rosa26-CreERT2*^*+*^ (*Setd4*^*flox/flox*^, blue filled cycle); 2) Tamoxifen-treated *Setd4*^*flox/flox*^;*Rosa26-CreERT2*^*+*^ (*Setd4*^*Δ/Δ*^, red square); and 3) tamoxifen-treated *Setd4*^*wt/wt*^;*Rosa26-CreERT2*^*+*^ (Tam-*Setd4*^*wt/wt*^, filled black cycles). The mice in groups 1 and 2 were male littermates. In all panels, each dot represents the data from an independent mouse, and none-irradiated mice are shown as day 0. *: p<0.05, **: P<0.01.

### Enhanced splenic erythropoiesis in *Setd4* deficient mice post TBI

In adult rodents, erythropoiesis in peripheral lymphoid organs occurs to support the animals when BM repopulation is impaired(45). During the course of the study, we noticed that *Setd4*^*Δ/Δ*^ mice had enlarged spleens more frequently than control mice (Fig. 3A-B). This prompted us to investigate whether there was any effect of *Setd4* loss on splenic erythropoiesis. Based on the stages of erythroid maturation (Fig. 3C), we analyzed cell surface markers CD71 and Ter119, which are commonly used to define the maturation stages of erythroid progenitor cells(46). We observed significantly more cells in the S3 (CD71^hi^/Ter119^hi^) stage and correspondingly reduced numbers of cells in the immature S1 (CD71^−^/Ter119^−^) stage at 2 weeks post TBI (Fig. 3D-E). There were significantly more late-stage cells (S2-S5) in the spleens of *Setd4*^*Δ/Δ*^ than in *Setd4*^*flox/flox*^ and *Setd4*^*wt/wt*^ mice. In addition, the percentage of erythroid cell population (S2-S5), relative to lymphoid and myeloid cell populations, were significantly increased in *Setd4*^*Δ/Δ*^ mice compared to the other groups in the spleens (Fig. 3F). These data indicate that loss of *Setd4* enhanced splenic erythropoiesis after TBI.

**Figure 3.**
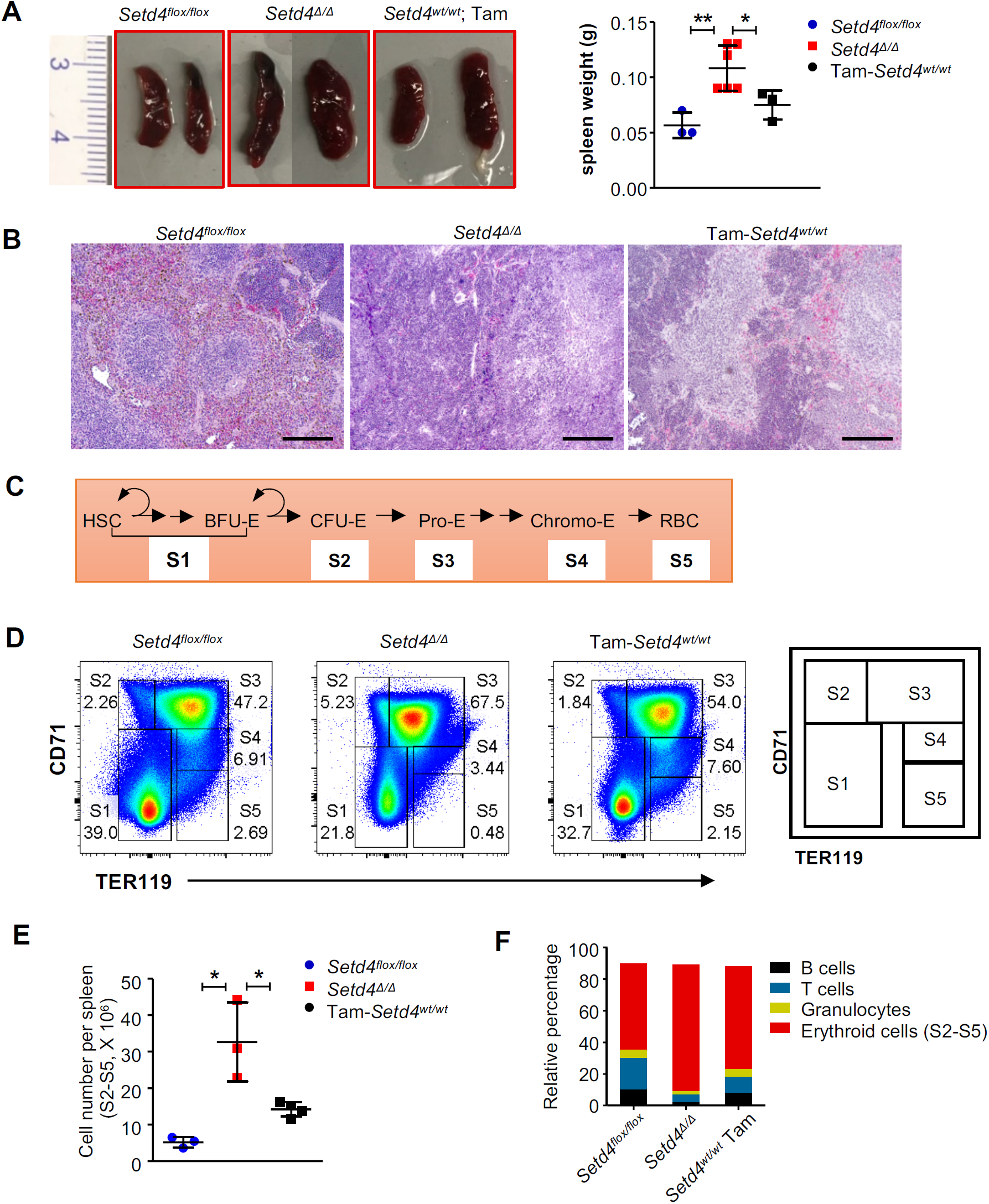
Enhanced splenic erythropoiesis in *Setd4* deficient mice after 8 Gy of TBI. The spleens were collected to analyze their cellular components. (A) Representative gross features of spleens, and spleen weight at 2 weeks post 8 Gy TBI from mice with the indicated genotypes. (B) Representative H&E staining of spleens at 2 weeks post 8 Gy TBI, scale bar: 100μm. A denser population of cells were observed in the *Setd4* deficient mice than the control mice. (C) A schematic representation of developmental sequence of erythroid cells from Stage 1 (S1) to Stage 5 (S5). HSC: hematopoietic stem cell; BFU-E: burst-forming unit-erythroid; CFU-E: colony-forming unit-erythroid; Pro-E: pro erythroblast; Chromo-E : chromatophilic erythroblasts; RBC: reticulocyte and erythrocyte. (D) Representative CD71 and TER119 staining profiling for spleen erythroid populations for the indicated mouse genotypes. The gating scheme to identify the different erythroid cell populations is shown on the right. (E) The numbers of total splenic erythroid cell populations (S2-S5) at 2 weeks post 8 Gy. (F) Percentage distribution of B cells, T cells, granulocytes, and erythroid cells in the spleens at 2 weeks post 8 Gy for the indicated mouse genotypes.

### *Setd4*^*Δ/Δ*^ reconstituted BM and engrafted *Setd4*^*Δ/Δ*^ HSCs were more sensitive to IR than wild type counterparts

It is generally believed that both the fitness of the stem and progenitor cells in the hematopoiesis hierarchy, and the regulatory function of the BM microenvironment niche contribute to the recovery from the hematopoietic failure(7,47). To understand the mechanism(s) by which *Setd4* deficiency conferred an enhanced BM recovery, we first investigated whether *Setd4* deletion confers radiation resistance to the stem and progenitor cells in the hematopoietic hierarchy using two independent approaches.

First, we measured the role of *Setd4* in the sensitivity of hematopoietic stem and progenitor cells (Fig. 4A). Bone marrows of the wild type B6.CD45.1 recipient mice were depleted by 2 doses of 6.5 Gy TBI separated by 4 hours, which is known to completely depletes BM and peripheral hematopoietic cells while sparing the BM niche(48). Within 24 hours, the recipient mice were transplanted with 1 million of Lin-cells isolated from the male littermates of *Setd4*^*Δ/Δ*^ or *Setd4*^*flox/flox*^, which express the CD45.2 congenic surface marker. Both the CD45 staining of peripheral blood cell from transplanted recipients and the genotyping of DNA extracted from ear clips 8 weeks after transplantation (Fig. 4B and 4C) confirmed that the donor HSCs had successfully implanted in the recipient mice and were functioning as the source of hematopoiesis. Thus, these mice were chimeric mice where the HSC-hierarchy was contributed by the donor HSCs (*Setd4*^*flox/flox*^ or *Setd4*^*Δ/Δ*^), but the BM niche and other stromal cells were derived from the recipient wild type mice.

**Figure 4.**
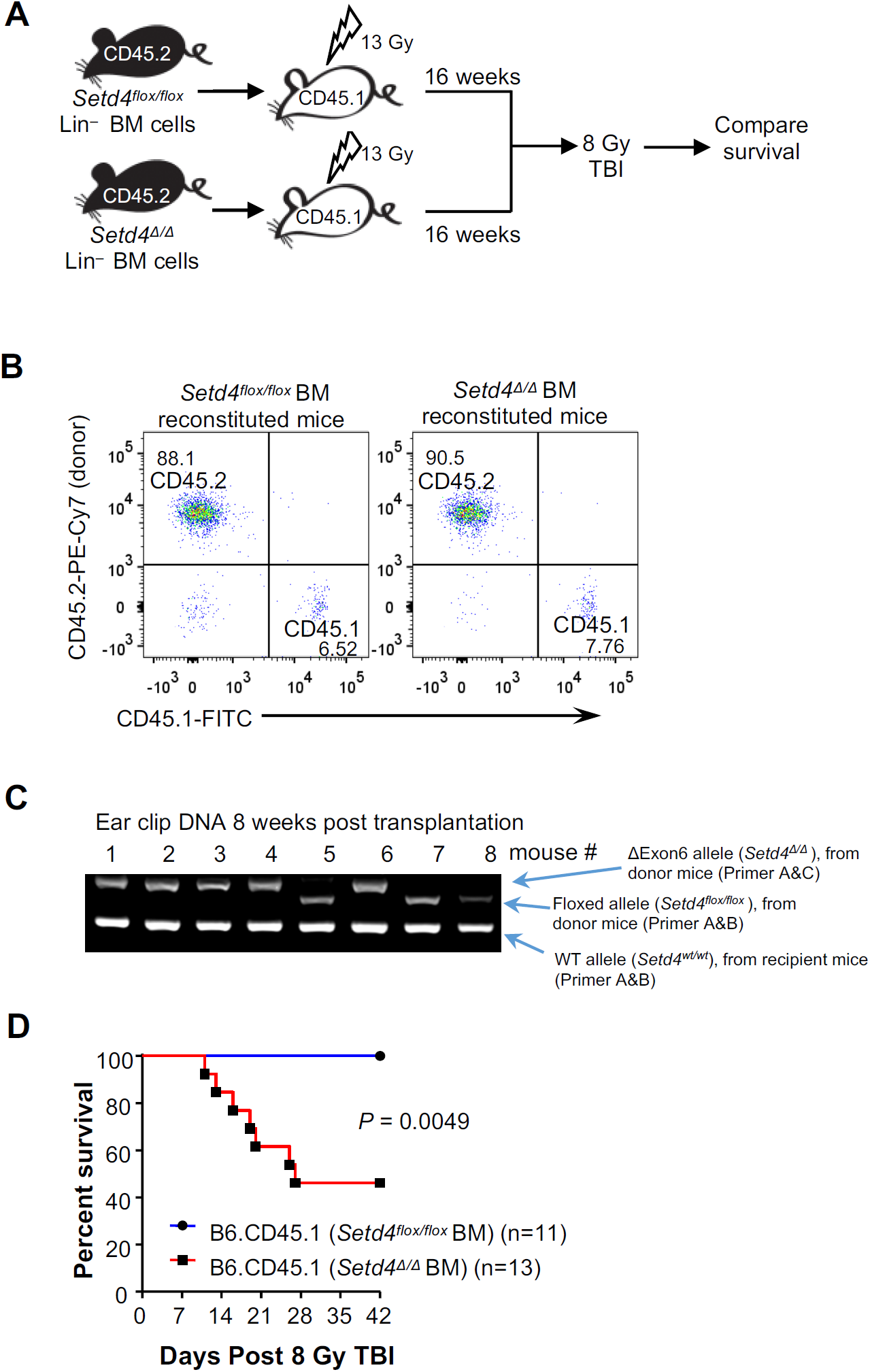
Radiation sensitivity of wild type and Setd4 deficient BM chimeric mice. (A) Schema outlining the non-competitive BM reconstitution assay. B6.CD45.1 recipient mice (B6 mice expressing the congenic CD45.1 marker) were exposed to 2 split doses of 6.5 Gy (total of 13 Gy) separated by 4 hours to deplete their native BM. Then one million Lin– BM cells from sex-matched littermates of either *Setd4*^*flox/flox*^ or *Setd4*^*Δ/Δ*^ mice (with CD45.2 marker) were transplanted into recipient mice. Eight weeks after the transplantation, peripheral blood and ear clips were sampled to confirm the successful implantation of the donor BM. Sixteen weeks after the transplantation, the mice were exposed to 8 Gy of TBI to induce hematopoietic syndrome and to address whether there is any difference of sensitivity caused by the transplanted donor BM. (B) Representative flow cytometric profiles of peripheral blood cells from the B6.CD45.1 recipients reconstituted with CD45.2 *Setd4*^*flox/flox*^ or CD45.2 *Setd4*^*Δ/Δ*^ BM cells. The peripheral blood cells were sampled 8 weeks after BM transplantation, and then stained antibodies to differentiate the transplanted cells (CD45.2+) from the recipient cells (CD45.1+). The small percentage (6-7%) of the remaining CD45.1 host cells are likely radio-resistant that were not killed by the irradiation. (C) Conformation of the genotyping of the BM chimeric mice. Genomic DNA were extracted from ear snips (containing blood) of recipients mice that were reconstituted with *Setd4*^*Δ/Δ*^ BM cells (lanes 1, 2, 3, 4, and 6) or with *Setd4*^*flox/flox*^ BM cells (lanes 5, 7, 8). Multiplex PCR genotyping was performed to simultaneously identify the wild type, floxed exon 6, and ΔExon6 alleles using three mixed primers: A (5’-TCCTGGGCTCTGCCATCCATG), B (5’-CTGTTGCAATGGAAATGCCAG), and C (5’-CTAAAGCTCTGCCCTAAGGTC). Pairing between primers A and B can amplify a 234bp wt allele and a 318bp flox-Exon6 allele but cannot amplify the ΔExon6 allele because region of primer B is deleted in the ΔExon6 allele. Paring between primers A and C can amplify a 369bp ΔExon6 allele. (Details of the PCR genotype strategies can be found in the Supplement Information of a previous report(40)). (D) Kaplan-Meier survival curves of BM chimeric mice after γ-radiation. Sixteen weeks after transplantation, the B6.CD45.1 recipients were treated with 8 Gy of TBI, and animal survival from the expected hemopoietic failure was monitored. Shown are the survival curves of the mice transplanted with *Setd4*^*Δ/Δ*^ (n=13) or *Setd4*^*flox/flox*^ (n=11). p = 0.0049.

To allow the transplanted LT-HSCs to replenish the entire hematopoiesis hierarchy(49), the recipient chimeric mice were kept for 16 weeks, and then treated with TBI (8Gy). Surprisingly, the mice transplanted with *Setd4*^*Δ/Δ*^ HSCs actually were more sensitive than the mice transplanted with the *Setd4*^*flox/flox*^ HSCs (Fig. 4D). This finding is contradictory to our initial prediction that the *Setd4* deletion may confer a resistance of the cells in the HSC hierarchy, and suggested that *Sedt4* deletion actually made the cells in the hematopoietic hierarchy more sensitive than *Setd4*^*flox/flox*^mice, which did not explain why the *Setd4*^*Δ/Δ*^ mice were more resistant to radiation induced hematopoietic syndrome than the *Setd4*^*flox/flox*^ mice.

To verify the above findings, we carried out a competitive transplantation assays as outlined in Fig. 5A. First, we depleted the BM of wild-type recipient mice (CD45.1) as described previously(48). Then, Lin-BM cells of *Setd4*^*Δ/Δ*^ (CD45.2) and *Setd4*^*wt/wt*^ (CD45.1/2) were mixed at approximately 1:1 ratio and transplanted into the recipients (CD45.1). The relative contribution of each type of donor cells to hematopoiesis was assessed by determining the ratio of CD45.2 over CD45.1/2 in peripheral blood cells with flow cytometric analysis at 4-week intervals, during the first 8-16 week after transplantation. As demonstrated in Fig. S5A, the donor *Setd4*^*Δ/Δ*^ (CD45.2) and competitor wild type (CD45.1/2) cells successfully implanted and there were few recipient (CD45.1) cells in the peripheral blood.

**Figure 5.**
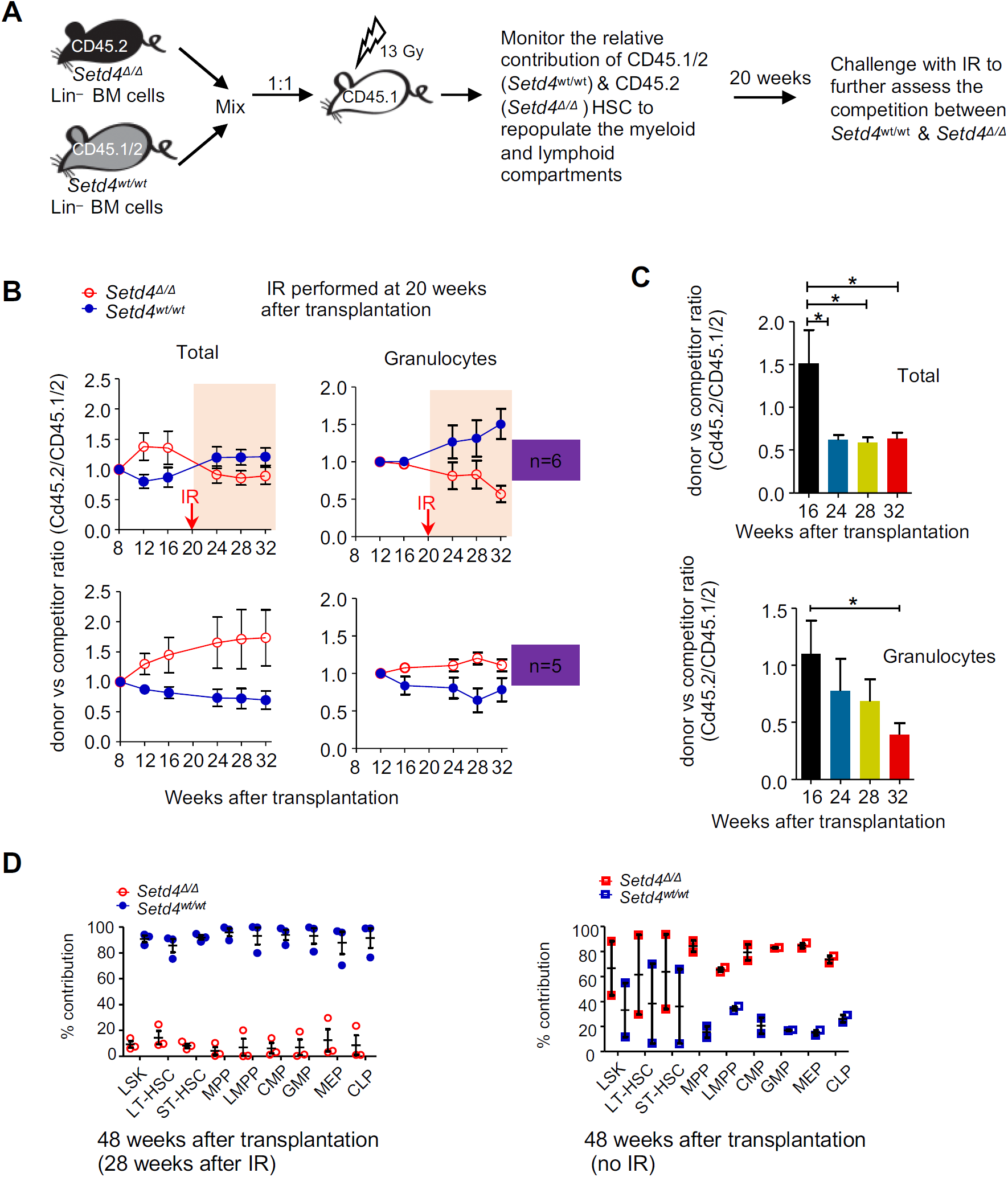
Radiation sensitivity of engrafted HSCs from *Setd4* deficient mice. (A) Shema detailing the competitive repopulation assay. Lin–BM cells (>95% Lin–) from *Setd4*^*Δ/Δ*^ mice (CD45.2) were mixed, at a ratio of 1:1, with WT competitor Lin–cells from B6.CD45.1/2 mice, and 1 × 10^6^ total cells were injected i.v. into lethally irradiated B6.CD45.1 recipients. The contribution of HSCs to reconstitute and subsequent refurbishment of the peripheral hematopoietic cells was monitored by flow cytometric analysis every 4 weeks. Twenty weeks after the transplantation, 6 of 11 reconstituted mice were exposed to sublethal irradiation (6.5 Gy) (IR: n=6; non-IR: n=5), and peripheral blood was monitored for 12 weeks, until final sacrifice for BM analysis at 18 weeks after the irradiation (or 48 weeks after the transplantation). (B) Peripheral blood from lethally irradiated transplanted mice were monitored for the relative contribution of *Setd4*^*Δ/Δ*^ donor (CD45.2) and wild type competitor (CD45.1/2) cells every 4 weeks, starting 8 weeks post-transplantation. Shown are the ratio of CD45.2 cells over CD45.1/2 competitor cells in the total peripheral blood cells and the granulocytes. Data show mean ± SEM among the mice, and are representative of two identical experiments. See Table S4 for the number of animals in each group. (C) The relative contribution of irradiated *Setd4*^*Δ/Δ*^ (donor) and WT (competitor) cells in all cells (left) or granulocytes (right) was plotted at 16, 24, 28 and 32 wk post transplantation. *, *p* < 0.05 by t-test. (D) The relative contribution of *Setd4*^*Δ/Δ*^ *and Setd4*^*wt/wt*^ BM HSCs and HPCs at 48 weeks after the transplantation. Left: irradiated mice (3 mice per group). Right: non-irradiated mice (2 mice per group). Mean and SEM are also shown. See Fig. S5 for representative flowcytometry profiles.

We reasoned that if the radiation sensitivities of *Setd4*^*Δ/Δ*^ (CD45.2) and *Setd4*^*wt/wt*^ (CD45.1/2) BM cells are similar, then the relative percentage of these populations should remain consistent in the mice after irradiation. The reconstituted mice were kept for 16 weeks and then exposed to 6.5 Gy of gamma-radiation 20 weeks after the initial transplantation. We then measured the relative population sizes of the *Setd4*^*Δ/Δ*^ (CD45.2) and *Setd4*^*wt/wt*^ (CD45.1/2) cells in peripheral blood every 4 weeks after the irradiation (i.e. 24, 28, and 32 weeks after the initial transplantation). Fig. 5B shows the relative population sizes of the irradiated (top panel) and non-irradiated (bottom panel) mice. When comparing the donor ratio at 16 weeks (4 weeks before irradiation) with that at 24 weeks (4 weeks after irradiation), a remarkable reduction in the CD45.2 (*Setd4*^*Δ/Δ*^) cells was observed (Fig. 5B, S5B), while there was an increase in these cells in non-irradiated mice (Fig. 5B & S5B). The total ratio of *Setd4*^*Δ/Δ*^ cells to wild type competitor cells dropped from 1.5:1 to 0.5:1 by 4 weeks after irradiation and never recovered at later time points (Fig. 5C). This indicated that upon irradiation the *Setd4*^*Δ/Δ*^ hematopoietic cells had reduced competence in the recipient mice when compared with the *Setd4* proficient cells. These results were in agreement with results observed above (Fig. 4) and confirmed that the *Setd4* deficient hematopoietic cells were more sensitive to radiation. Another important finding from this experiment is that, in the absences of irradiation, the *Setd4* deficient hematopoietic cells may have had an advantage in re-populating the BM over the Setd4^wt/wt^ hematopoietic cells (see bottom panels of Fig. 5B, S5B).

To assess the long-term fitness of the BM cells, the above mice were sacrificed 28 weeks after irradiation (or 48 weeks after the initial transplantation) and lineage analysis was performed on their BM cells as described in Fig. 2 and S4. As shown in Fig. 5D and S6 the ratio of the *Setd4*^*Δ/Δ*^ to *Setd4*^*wt/wt*^ cells in the BM were significantly reduced at this late stage after the irradiation. These data again suggested that the *Setd4* deficient LT-HSCs and progeny were more sensitive to radiation. However, among the two available pairs of non-irradiated mice, there were significantly more early progenitors in *Setd4*^*Δ/Δ*^ than *Setd4*^*wt/wt*^, although the advantage of *Setd4*^*Δ/Δ*^ LSK, LT-HSC and ST-HSC was marginal. These data again implied that the *Setd4*^*Δ/Δ*^ cells had an advantage in repopulating the BM when not stressed by irradiation.

### *Setd4* deletion confers an improved BM microenvironment to support HSC transplantation

Collectively, data from the BM transplantation assays (Fig. 4 and 5) implied that *Setd4* deletion makes the hematopoietic cells acutely more sensitive to radiation, while conferring an advantage to the *Setd4*^*Δ/Δ*^ hematopoietic progenitors in the absences of irradiation. Nevertheless, these findings suggested that the improved recovery of *Setd4*^*Δ/Δ*^ mice from acute BM failure wasn’t due to a radiation resistance of the *Setd4*^*Δ/Δ*^ HSC or HPC at the time of exposure. This led us to form an alternative hypothesis that the enhanced recovery of *Setd4*^*Δ/Δ*^ mice from acute hematopoietic syndrome may be attributed to a more favorable *Sedt4* deficient BM niche. To test this, we performed HSC dilution survival assays (Fig. 6A). Male *Setd4*^*flox/flox*^:*Rosa26-CreERT2* littermates were injected with Tam (*Setd4*^*Δ/Δ*^) or Oil (*Setd4*^*flox/flox*^). Seven days later, the mice were subjected to lethal doses of radiation (2 doses of 6.5 Gy separated by 4 hours) to deplete the BM, and then transplanted with titrated numbers of wild-type Lin^−^ BM cells from CD45.1 mice (Table S5). In this assay, the survival of recipients depends on whether the damaged BM niche is able to support transplanted HSCs after limiting numbers are transplanted. As shown in the Fig. 6B, *Setd4*^*Δ/Δ*^ had a significantly improved survival than *Setd4*^*flox/flox*^ when transplanted with the same number of wild type Lin-cells. Based on a survival calculation, the number of cells required for 63% survival (or 37% fatality rate) was 43.8 × 10^3^ for *Setd4*^*Δ/Δ*^ mice and 69 ×10^3^ for *Setd4*^*flox/flox*^ mice, a reduction of 1.6-fold. This indicates that *Setd4*^*Δ/Δ*^ mice have a more favorable BM environment to host the HSC transplantation. Thus, the radiation-resistance of *Setd4* deficient mice was likely due to an improved BM niche that nourishes the damaged hematopoietic cells. This highlights the importance of the BM niche in the recovery of the hematopoiesis in contrast to the intrinsic radiation sensitivity of the hematopoietic cells themselves.

**Figure 6.**
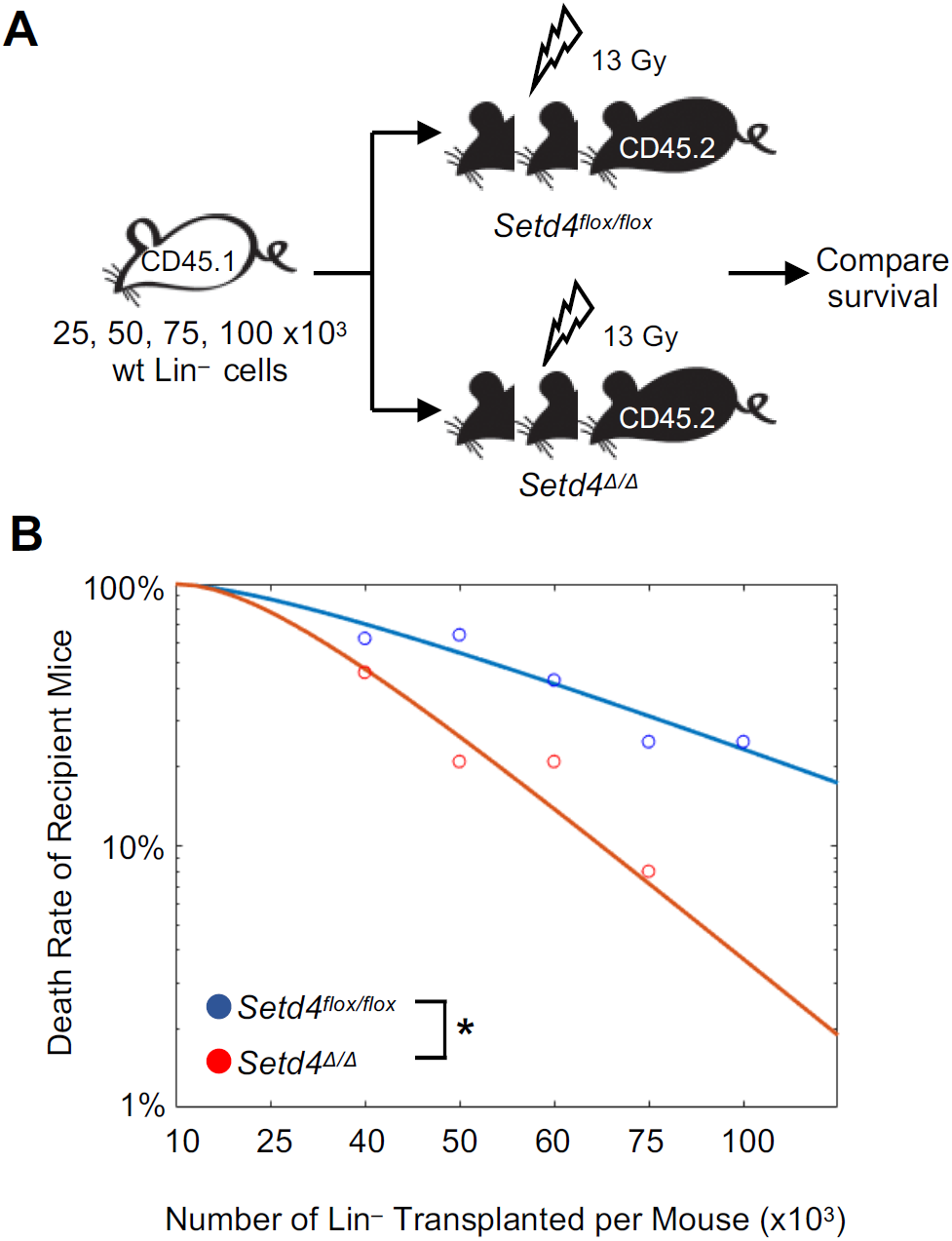
Survival of lethally-irradiated *Setd4* deficient and proficient mice after transplantation of wild type Lin-BM donor cells. (A) Shema detailing HSC transplantation to rescue the lethally irradiated recipient mice. Serially diluted donor Lin-BM cells (10, 25, 40, 50, 60, 75, and 100×10^3^) from B6.CD45.1 mice were transferred into lethally irradiated *Setd4*^*Δ/Δ*^ or *Setd4*^*flox/flox*^ recipients. Survival was monitored for 30 days. (B) Death rate of the transplanted recipient mice as a function of the numbers of transplanted Lin^−^ cells per mouse. For each experimental point, 6-14 recipients were used (see Table S5 for the specific number of mice used in each time point). The curves were fit by nonlinear regression. The calculated number of cells for a 63% of survival rate is 69.0×10^3^ for *Setd4*^*flox/flox*^ mice, and 43.8×10^3^ for *Setd4*^*Δ/Δ*^ mice. P=0.029 by two-way Anova test.

## Discussion

In this study, we developed the first *Setd4*-knockout mouse model to directly identify the function of SETD4 in mammalians, and surprisingly found that *Setd4*-deficient mice were more resistant to radiation-induced hematopoietic syndrome. Interestingly, *Setd4* deletion conferred a radio-sensitivity to the HSCs *in vivo*, but the non-irradiated *Setd4* deficient HSCs had an advantage over wild type HSCs in recipient mice, and the BM niches of the *Setd4*-deficient recipient mice had an enhanced ability to engraft transplanted HSCs. Furthermore, there was an enhanced splenic erythropoiesis in *Setd4* deficient mice. These novel findings suggest a unique mode of action by which Setd4 modulates the recovery from BM failure and underscores the importance of the BM niche in the recovery from hematopoietic failure.

Given that BM injury is a major adverse side effect of anticancer therapy and excessive exposure to radiation, it is possible that inhibition of Setd4 during these circumstances may be beneficial to the recovery of the BM. Based on our findings, it is foreseeable that inhibition of Setd4 would: 1) sensitize cells of the HSC lineage in the host to BM depletion, 2) make the host BM niche more amenable to receive allogeneic transplantation, and 3) improve the competence of the donor HSCs. Thus, Setd4 would potentially be an attractive target for inhibition to help condition the BM before allogeneic transplantation, a strategy used to treat widely disseminated malignancies and some solid tumors(8-10).

It is well known that the overall function of the hematopoietic system is both intrinsically regulated by the HSCs and their progenies, as well as by the BM niche nurturing the HSCs. One striking finding of our study is that the *Setd4* deficient HSCs are actually more sensitive to irradiation, but the BM niche of the same *Setd4* deficient mice is more receptive to the engraftment of HSC upon transplantation. However, the overall consequence of these confounding effects was an improved recovery from radiation induced hematopoietic failure. This finding underscores the importance of the BM niche in the overall function of the hematopoietic system.

It should be noted that our results also suggested that the improved recovery of *Setd4* deficient mice is independent of *p53* deletion, and *Setd4* loss further enhanced the survival of *p53*-deficient mice post IR (Fig. S1). Pharmacological inhibition of p53 has been proposed to protect individuals from acute BM injury and attenuate the development of radiation-related blood cancer(50). Similarly, it is foreseeable that targeting Setd4 may represent another strategy to protect BM from radiation and radiation-induced lymphomagenesis.

The mammalian SETD4 is largely an orphan methyl-transferase since no physiological substrate of mammalian SETD4 has been reported. Several preliminary reports have suggested a potentially oncogenic activity of mammalian SETD4(37-39), and oncogenic fusions of SETD4 had been reported in TCGA database. The conditional *Setd4* deletion did not significantly change spontaneous tumor formation(40), which is in an agreement with these preliminary reports. Clearly, this study has only just begun to reveal the function of mammalian SETD4, and additional studies would need to be performed to further understand SETD4 activities in mammalians.

In summary, our studies revealed a critical role of Setd4 in maintaining hematopoiesis, suggesting that *Setd4* inhibition might be an attractive strategy to mitigate radiation-induced BM injury, and improve the outcome of allogeneic HSC transplantation after TBI for patients with widely disseminated malignancies. Future studies will be needed to identify physiological SETD4 substrates and to elucidate the regulatory mechanisms of SETD4 expression.

## Acknowledgments

This research was supported by NIH R01CA156706 and R01CA195612 grants to ZS and R21AI105747 to LKD, by support from Robert Wood Johnson Foundation to CINJ, and by the Biospecimen Repository and Histopathology Service, Genome Editing, and Flow Cytometry/Cell Sorting shared resources of The Rutgers Cancer Institute of New Jersey (P30CA072720). The authors thank Ms. Neta Schneider for assistance with some of the mouse work, Mr. Louis Osorio (Rutgers Child Health Institute of New Jersey) for generous technical assistance with performing the mouse BM transplantation studies; and Dr. Arthur Roberts (Rutgers Cancer Institute of New Jersey) for cell sorting; Dr. Ping Xie and Dr. Sam Bunting (Rutgers the State University of New Jersey) for suggestions throughout the study.

## Authorship Contributions

XF and ZS: designed the experiments, performed data analysis, and drafted the initial versions of manuscript. XF: performed most of the experiments; HL, JL, and MS: performed and assisted with some of the experiments. HL, and JY: performed some of the early studies led to the construction of the mouse models; LD, and SD: provided resources for some experiments, computational analysis, and edited the manuscript. ZS: directed and oversaw the project, secured funding for the study, completed the final version of the manuscript.

