## Supplementary figures and images for "Deletion of mouse *Setd4* promotes the recovery of hematopoietic failure"

### Supplement Materials and Methods, Tables, and Figures

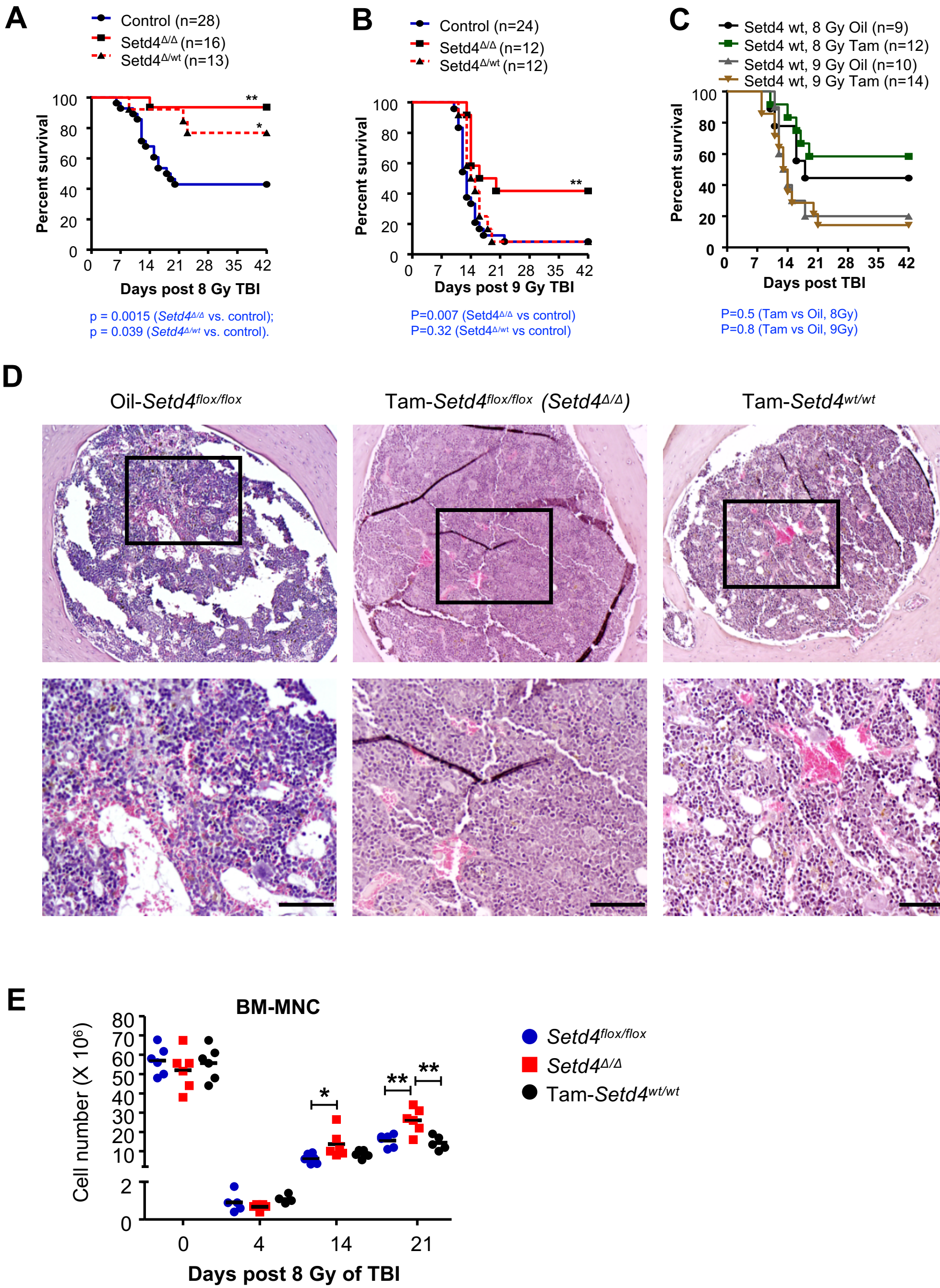

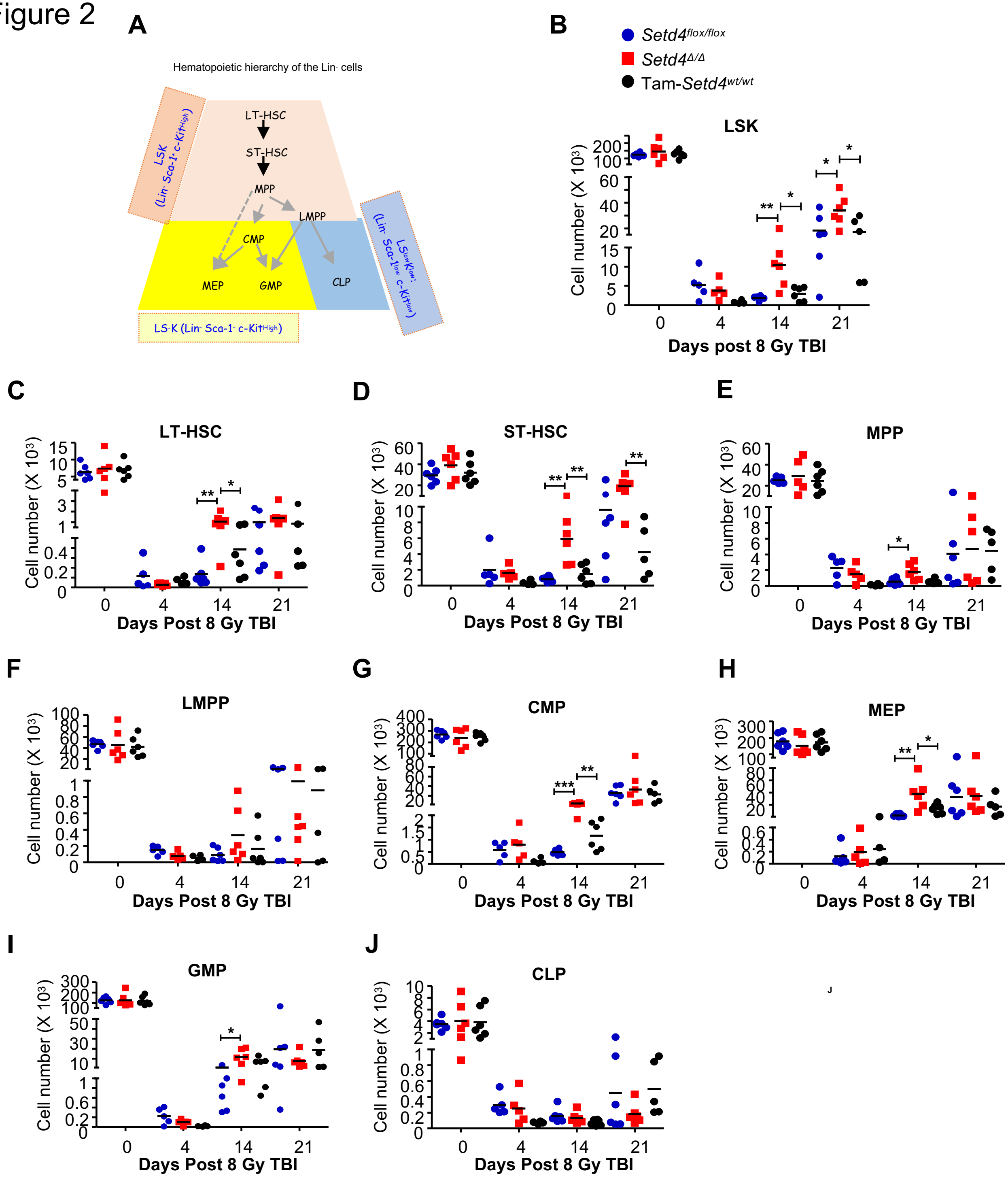

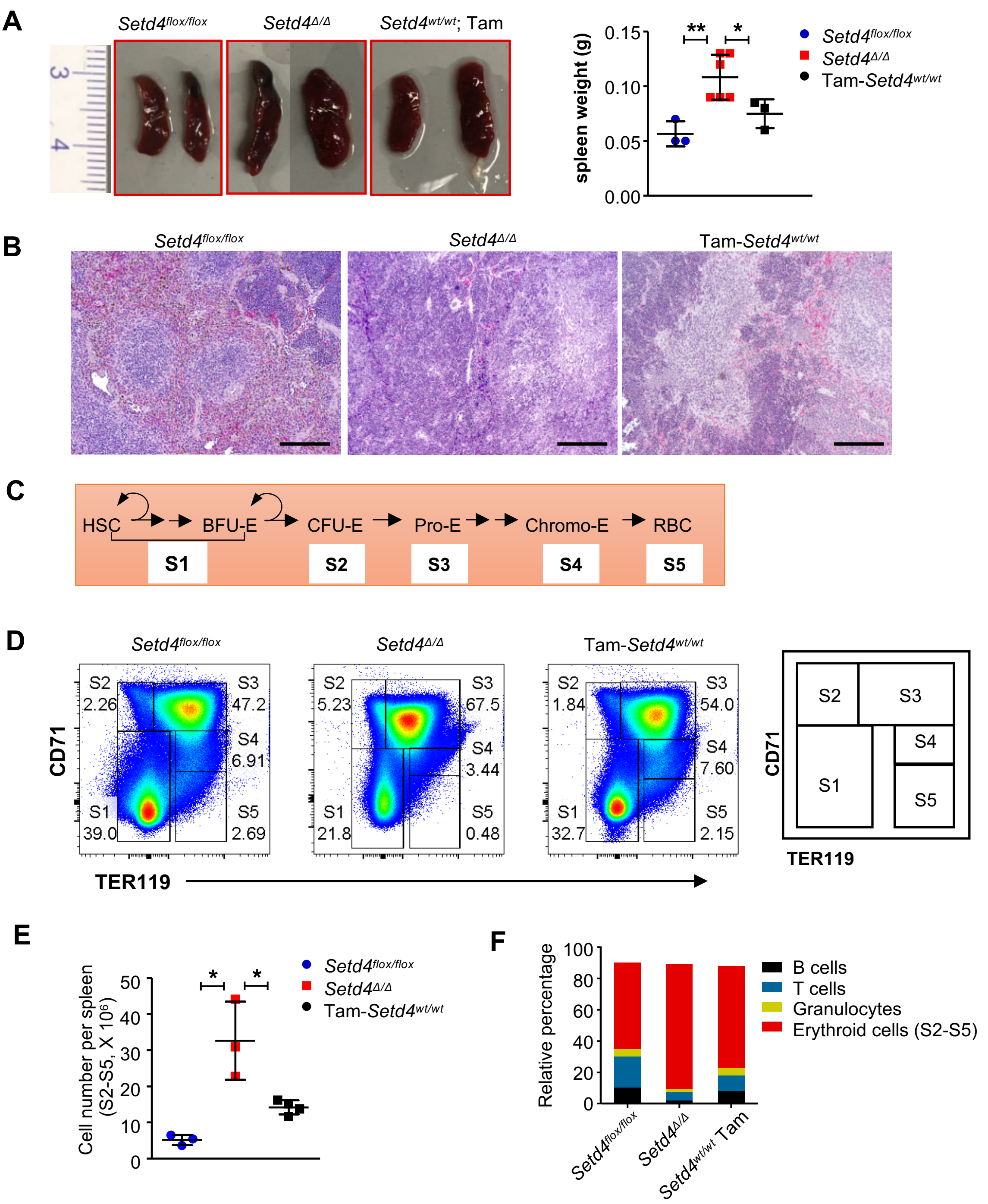

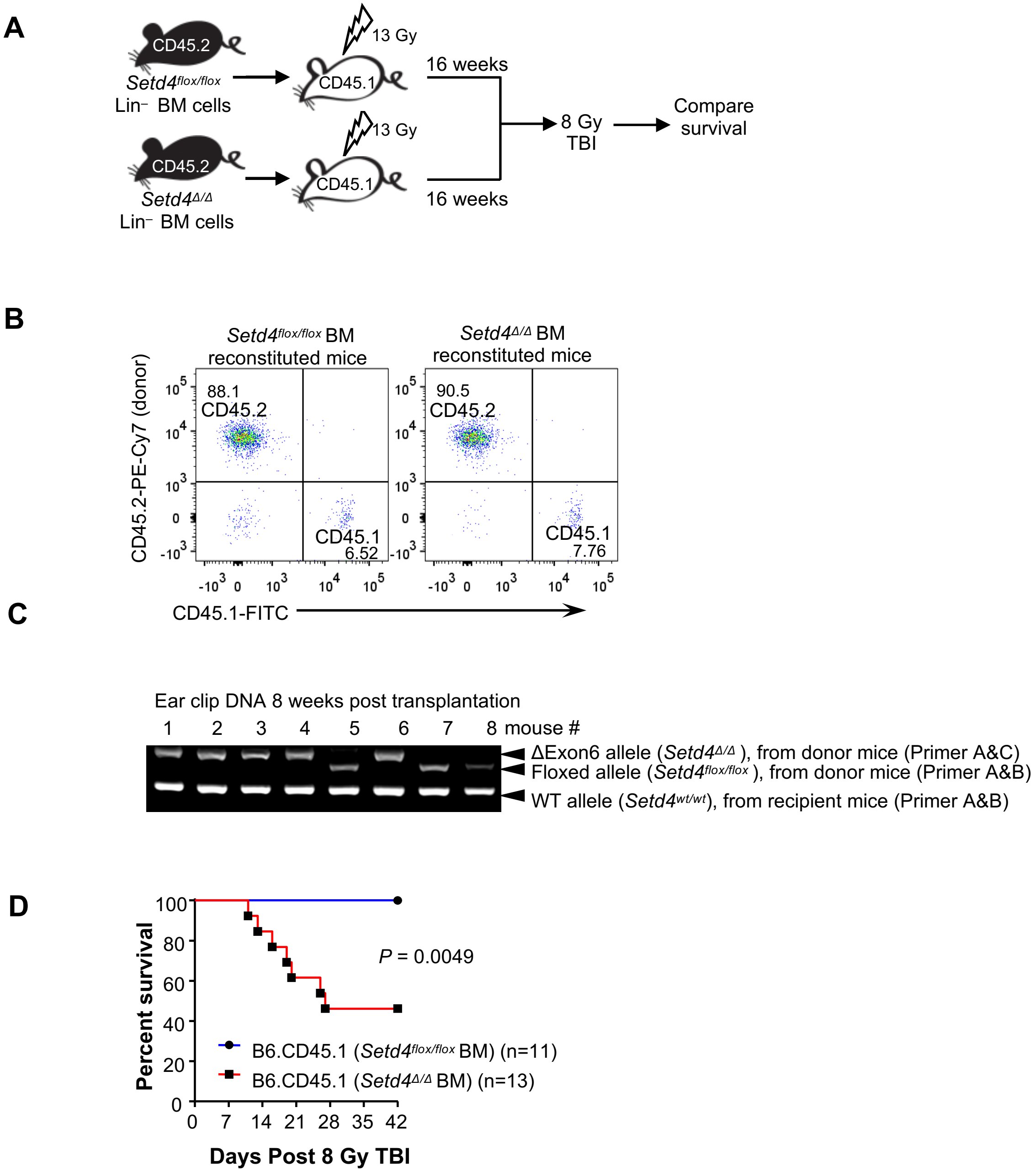

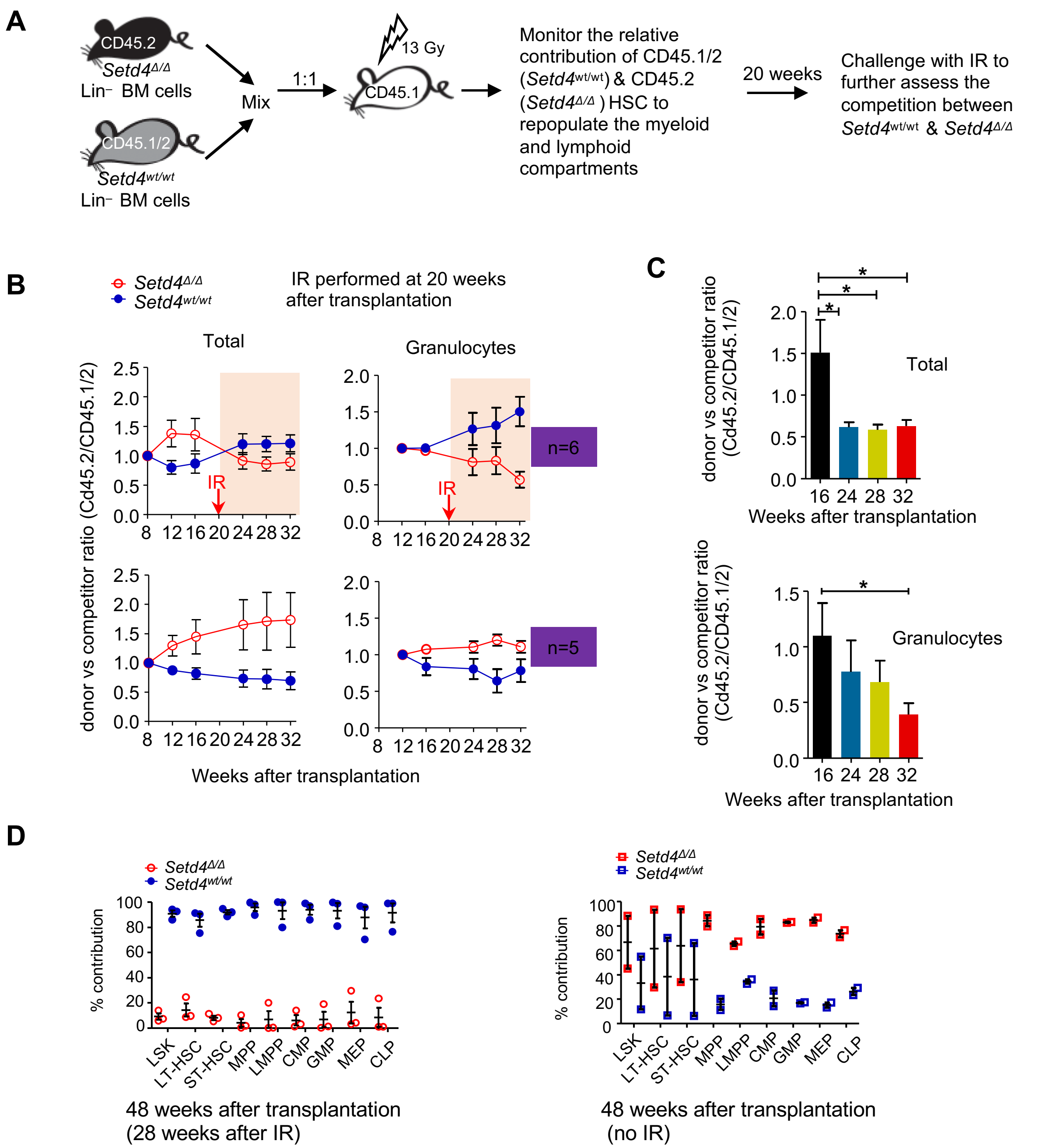

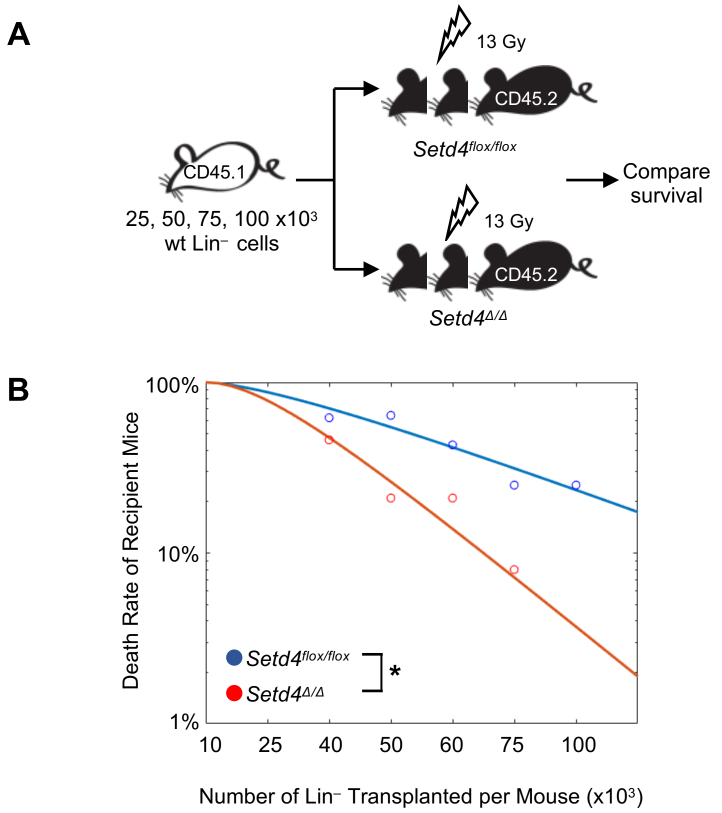
